# A Msp1-containing complex removes orphaned proteins in the mitochondrial outer membrane of trypanosomes

**DOI:** 10.1101/2023.03.07.531462

**Authors:** Markus Gerber, Ida Suppanz, Silke Oeljeklaus, Moritz Niemann, Sandro Käser, Bettina Warscheid, André Schneider, Caroline E. Dewar

## Abstract

The AAA-ATPase Msp1 extracts mislocalized outer membrane proteins and thus contributes to mitochondrial proteostasis. Using pull down experiments we show that trypanosomal Msp1 localizes to both glycosomes and the mitochondrial outer membrane, where it forms a stable complex with four outer membrane proteins. The trypanosome-specific pATOM36 mediates complex assembly of a-helically anchored mitochondrial outer membrane proteins such as protein translocase subunits. Inhibition of their assembly triggers a pathway that results in the proteasomal digestion of unassembled substrates. Using inducible single, double and triple RNAi cell lines combined with proteomic analyses we demonstrate that not only Msp1 but also the trypanosomal homolog of the AAA-ATPase VCP are implicated in this quality control pathway. Moreover, in the absence of VCP three out of the four Msp1-interacting mitochondrial proteins are required for efficient proteasomal digestion of pATOM36 substrates suggesting they act in concert with Msp1. pATOM36 is a functional analogue of the yeast MIM complex and possibly of human MTCH2 suggesting that similar mitochondrial quality control pathways linked to Msp1 might also exist in yeast and humans.

## Introduction

The mitochondrial outer membrane (OM) forms the interface between mitochondria and the cytosol, and many of its proteins have important functions in cytoplasmic-mitochondrial communication. Integral OM proteins are often α-helically anchored, and many have just a single transmembrane domain (TMD). Intriguingly, there are at least three unrelated protein factors mediating the biogenesis of α-helically anchored OM proteins in different eukaryotic clades. The mitochondrial import complex (MIM), was discovered in *Saccharomyces cerevisiae* and consists of two small proteins, Mim1 and Mim2, that are restricted to fungi (1–4). In the parasitic protozoan *Trypanosoma brucei*, the kinetoplastid-specific peripheral atypical translocase of the outer membrane 36 (pATOM36) has the same function (5, 6). Expression of pATOM36 in yeast lacking the MIM complex restores growth under non-permissive conditions, and vice versa, expression of the MIM complex complements the OM protein biogenesis defect in pATOM36-ablated trypanosomes (7). In human cells, the mitochondrial animal-specific carrier homolog 2 (MTCH2) is necessary and sufficient to insert α-helically anchored membrane proteins into the OM (8). However, at least in yeast, spontaneous insertion into the OM also seems possible for some proteins (9, 10).

Safeguarding of mitochondrial functions requires mitochondria-associated degradation (MAD) pathways that survey the OM and guarantee that its proteins are correctly targeted and assembled (11–13). The highly conserved Msp1, an ATPase associated with diverse cellular activities (AAA), plays a key role in in this process. It consists of an N-terminal TMD and a C-terminal AAA domain, which faces the cytosol (14), and localises to both the OM and the peroxisomal membrane. Msp1 extracts mislocalised and/or misassembled proteins from the OM and feeds them to the cytosolic proteasome (15–19). Non-mitochondrial tail-anchored (TA) proteins, which have a single TMD at their C-terminus, can be prone to OM mistargeting under both normal and stress conditions (15–18, 20–22). The latter includes a deficient guided-entry of TA proteins (GET) pathway in the ER or an impaired peroxisomal targeting machinery (15, 16, 23, 24). There is no clear sequence consensus between Msp1 substrates (15–18, 25, 26). However, Msp1 recognizes and extracts orphan TA proteins that are normally found in a complex, which suggests that their oligomeric state is an important determinant (18, 26). Intriguingly, mitochondrial Msp1 is not known to form stable complexes with other proteins and appears to extract its substrates from the OM without help from other proteins (17, 26). However, adaptor proteins may still be required for substrate selectivity or regulation of activity. For example Msp1 is able to clear stuck precursor proteins from the TOM complex via a transient interaction with the inducible peripheral OM protein Cis1 and the TOM receptor Tom70 in response to mitochondrial protein import stress (19).

Msp1 deletion in yeast causes a mild growth phenotype only, which suggests some redundancy in OM quality control (16, 27). In line with this, it was shown that under stress conditions the AAA-ATPase VCP, a soluble cytoplasmic component of the endoplasmic reticulum-associated protein degradation (ERAD) system, can also extract mistargeted proteins from the OM (28). For degradation by the proteasome, proteins generally require ubiquitination. It has been shown that mislocalised proteins can be extracted from the OM by Msp1 and transferred to the ER, where they are ubiquitinated by the ER-resident E3 ligase Doa10. This allows for their extraction from the membrane by VCP and subsequent degradation by the proteasome (26, 29). However, as E3 ligases normally have specific sets of substrates, this pathway might not be required for all Msp1 substrates, and some may be degraded by the proteasome without prior ubiquitination (29).

Studies of mitochondrial processes, including MAD pathways, have mainly focused on yeast and mammals, which belong to the same eukaryotic supergroup of the Opisthokonts. However, a better understanding of their basic features and evolutionary history requires that these processes are studied across divergent eukaryotes. Arguably the best studied mitochondrion outside of yeast and mammals is that of *Trypanosoma brucei*. It belongs to the Discoba supergroup, which is essentially unrelated to the Opisthokonts (30–32).

Here, we show that TbMsp1 is localized to both the mitochondrial OM and glycosomes, a kinetoplastid-specific organelle related to peroxisomes. Cells depleted of pATOM36 indicate that TbMsp1 removes destabilised OM proteins in concert with cytosolic TbVCP by delivering them to the proteasome. We found that four integral OM proteins stably interact with TbMsp1, and show that, uniquely, three of these interaction partners are required for the full activity of TbMsp1 in this quality control pathway, setting TbMsp1 apart from Opisthokont Msp1.

## Results

### TbMsp1 interacts with proteins of the mitochondrial OM and the glycosomes

Msp1 is highly conserved within eukaryotes, with TbMsp1 showing 34.5% and 33.5% identity to that of yeast and human Msp1 respectively. This conservation is in contrast to many other trypanosomal OM proteins, most of which are specific to kinetoplastids (33). TbMsp1 has the expected conserved sequence motifs including the AAA-domain and the Walker A and Walker B motifs required for ATP binding and hydrolysis (Fig. 1A, Fig. S1). To identify TbMsp1 interaction partners and determine its intracellular localisation, we produced a cell line expressing a C-terminally in situ HA-tagged TbMsp1 variant. Digitonin-extracted crude mitochondrial fractions of this cell line were subjected to a stable isotope labelling by amino acids in cell culture (SILAC)-immunoprecipitation experiment using anti-HA antibodies. TbMsp1-HA precipitated ten proteins with enrichment factors of more than 3-fold (Fig. 1B). From previous proteomic analyses, three were identified as OM proteins, five are glycosomal proteins, TbTsc13 showed both localisations, and Tb927.3.4500 is the cytosolic fumarate hydratase, which was hypothesised to interact with the cytosolic side of the glycosomal membrane (33–36). Of the glycosomal proteins, the peroxisome biogenesis protein Pex11 (Tb927.11.11520), tyrosine phosphatase (Tb927.10.10610), glycosomal metabolite transporters GAT1 (Tb927.4.4050) and GAT2 (Tb927.11.3130) all contain TMDs (37–39), whereas phosphoglycerate kinase A (Tb927.1.720) is localised in the glycosomal lumen (40, 41).

**Figure 1.**
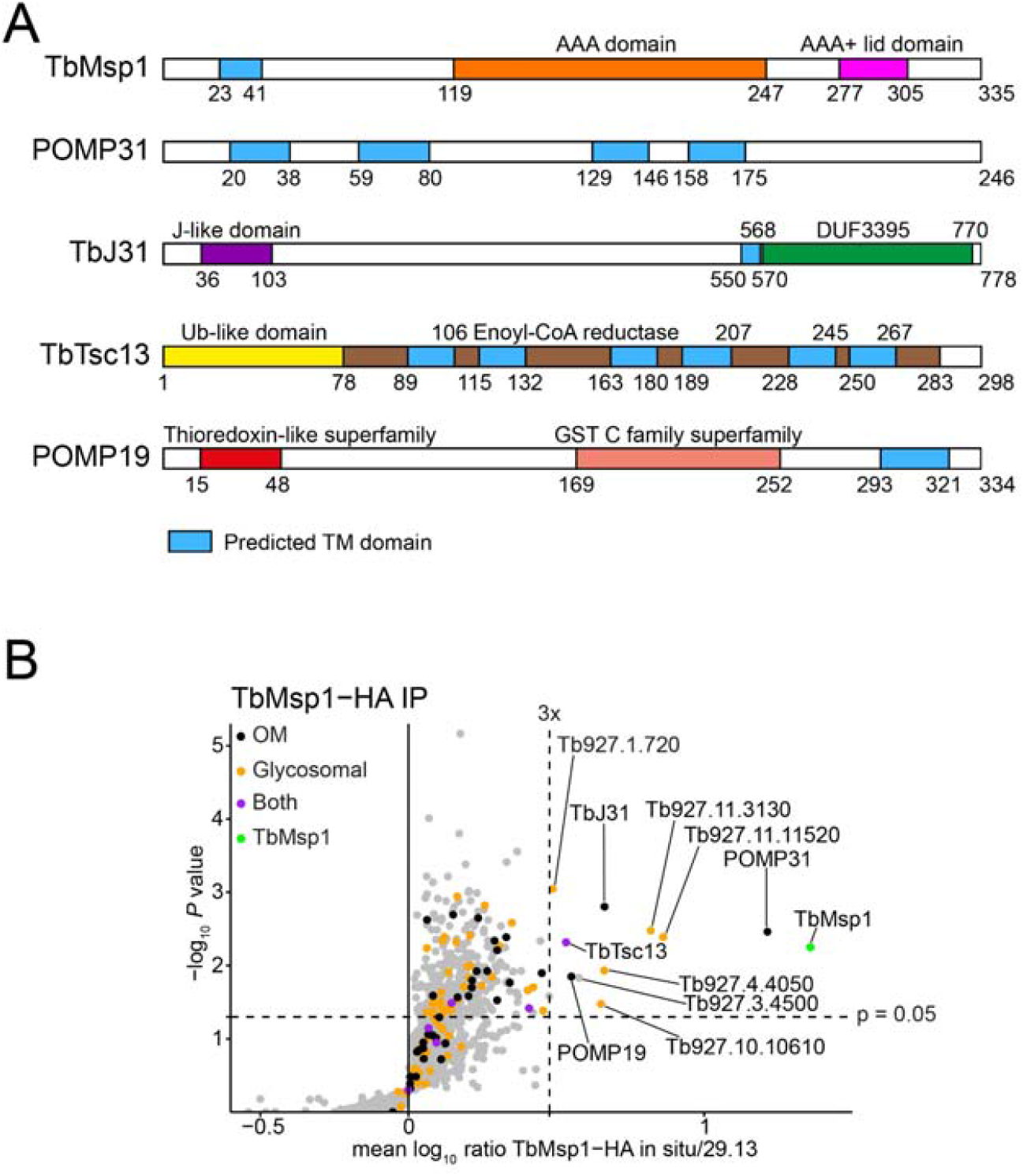
TbMsp1 forms complexes in the OM and glycosomes. A. Schematic depiction of predicted domain structures of TbMsp1 and four interacting OM proteins. The indicated domains were predicted as described in the material and methods section. B.TbMsp1 complexes were immunoprecipitated from crude mitochondrial fractions of differentially SILAC labelled 29.13 parent cells and cells expressing in situ tagged TbMsp1-HA and analysed by quantitative mass spectrometry (n=3). Proteins found to be significantly enriched more than 3-fold in TbMsp1 complexes are labelled with either their name or their accession number.

In the present study, we focused on the four most enriched OM proteins (Fig. 1A). The first is the protein of the mitochondrial OM proteome 31 (POMP31), which is a kinetoplastid-specific protein of unknown function with four predicted TMDs. The second one is TbJ31, a J-like protein which has a single predicted TMD (42). It is the homologue of mammalian DNAJC11, with which it also shares the domain of unknown function (DUF) 3395 (43). The third is POMP19, a kinetoplastid-specific protein with a single TMD that contains a predicted thioredoxin-like and a predicted glutathione S-transferase domain and the fourth is TbTsc13, which is also found in glycosomes. It shows homology to the mammalian enoyl-CoA reductase of the ER elongase complex and has six predicted TMDs (44). TbTsc13 contains a predicted ubiquitin-like domain at the N-terminus.

Subcellular fractionation of cells expressing TbMsp1-HA and its four epitope-tagged OM interactors showed that they all co-fractionate with the voltage dependent anion channel (VDAC), as would be expected for mitochondrial OM proteins (Fig. S2A). Moreover, all proteins were predominantly recovered in the pellet when subjected to carbonate extraction at high pH, indicating that, in line with their predicted TMDs, they are all integral membrane proteins.

Immunofluorescence of cells expressing either TbMsp1-myc or an epitope-tagged interactor revealed a close degree of co-localisation of POMP31, TbJ31 and POMP19 with the mitochondrial marker atypical translocase of the OM 40 (ATOM40) (Fig. S2B). Additionally, normalised abundance profiles of untagged native TbMsp1 and its four interactors from a previous proteomic analysis with six subcellular fractions, including crude and pure OM, confirm the OM localisation of all four proteins (Fig. S2C) (33).

As expected from the SILAC-pulldown experiment (Fig. 1B) and previous analyses (34, 44), TbMsp1-myc and TbTsc13-HA are not exclusively mitochondrially localised. TbMsp1-myc, in addition to mitochondrial staining, partially co-localised with the glycosomal marker aldolase (ALD) (Fig. S3). The localisation of TbTsc13-HA, in line with its predicted function as an enoyl-CoA reductase, partially overlapped with the ER luminal binding protein (BiP) (Fig. S3).

Next, we validated the interactions of the four proteins with TbMsp1, and between each other, by immunoprecipitations using cell lines in which both Msp1 and one candidate interactor were epitope-tagged. Interactions could be confirmed between TbMsp1 and each of POMP31, TbJ31, POMP19 and TbTsc13 (Fig. 2A), whereas interactions were not detected between these proteins and the most abundant OM protein, VDAC. Using the same method, we could also detect mostly reciprocal interactions between POMP31, TbJ31, POMP19 and TbTsc13. These results suggest that all five proteins are present in the same stable protein complex (Fig. 2B).

**Figure 2.**
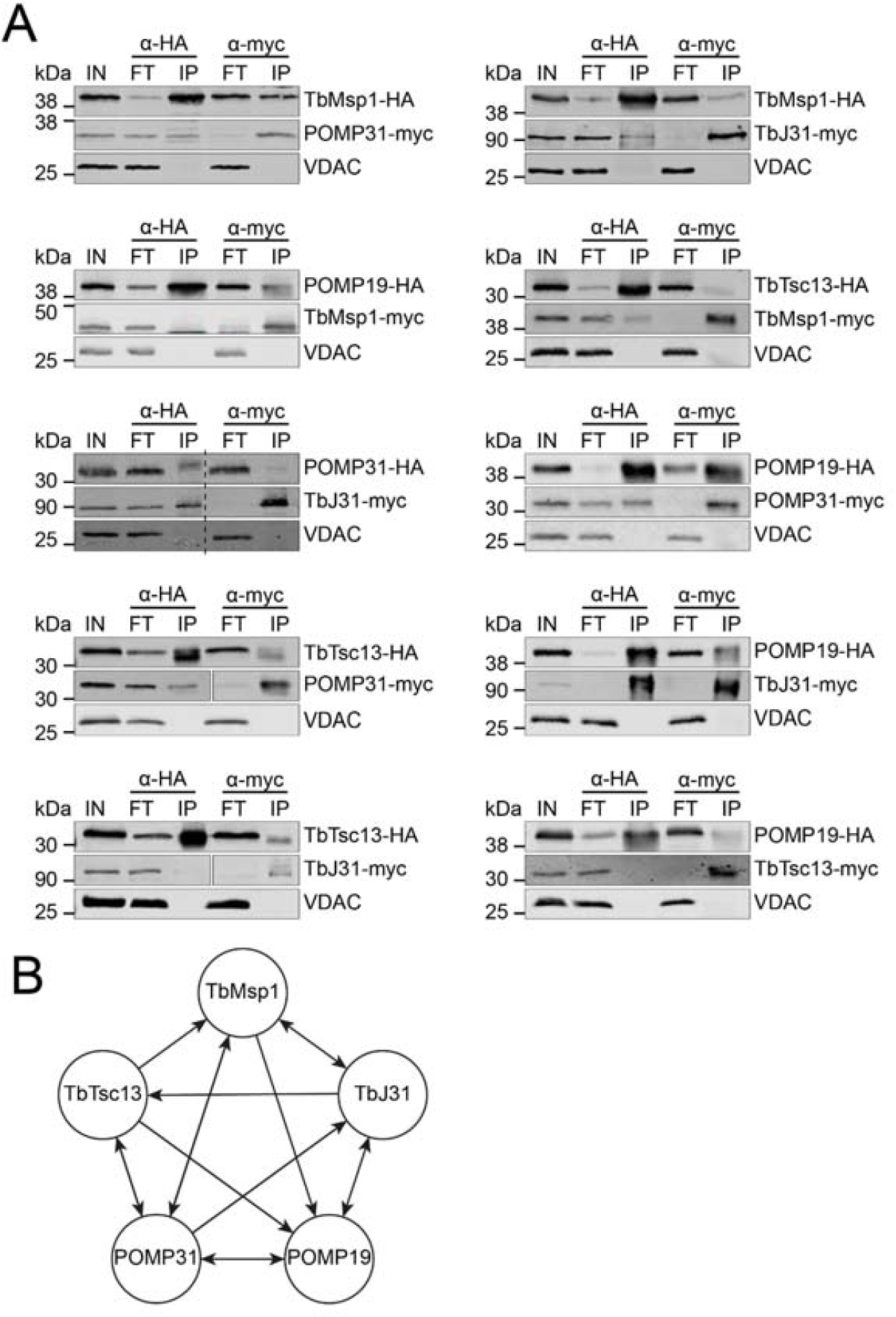
Reciprocal IPs reveal a TbMsp1-centric interaction network in the OM. A. Crude mitochondrial fractions from cells overexpressing the indicated C-terminally myc- and HA-tagged proteins were analysed by immunoprecipitation. Crude mitochondrial fractions (IN), unbound proteins (FT), and final eluates (IP) were separated by SDS-PAGE. Resulting immunoblots were probed with anti-tag antibodies and antisera against VDAC. B.Summary of the confirmed interactions detected by coimmunoprecipitation. Two sided arrows indicate reciprocal interactions.

Finally, to investigate the importance of TbMsp1 and the four TbMsp1-interacting proteins for cell viability, we produced inducible RNAi cell lines targeting the ORFs of these proteins. However, despite the fact that the RNAi efficiently depleted the corresponding target mRNAs (Fig. S4, insets), only the RNAi cell line targeting TbTsc13 showed a clear inhibition of growth (Fig. S4, bottom panel). This was expected as TbTsc13 is likely to play an essential role in fatty acid elongation as in yeast (45). Thus, within the limit of the RNAi analysis, which does not completely deplete gene products, TbMsp1, POMP19, POMP31 and TbJ31 are not essential for normal cell growth in the procyclic form of trypanosomes.

### pATOM36-RNAi results in proteasomal depletion of its substrates

The biogenesis of many α-helically membrane-anchored mitochondrial OM proteins is mediated by distinct protein factors in yeast (MIM complex), humans (MTCH2) and trypanosomes (pATOM36) (1–6, 8). Moreover, for the MIM complex and pATOM36, reciprocal complementation experiments demonstrate that they are functionally interchangeable (7). In the present study, we focussed on the proteomic consequences of pATOM36 depletion in trypanosomes. The total cellular levels of the ATOM complex subunits ATOM19 and ATOM46 were massively reduced after induction of pATOM36 RNAi (Fig. 3A, lanes 1 and 2; Fig. 3B), in agreement with a previous proteomic analysis of crude mitochondrial fractions of pATOM36-depleted cells (6). This was confirmed when whole cell samples of the same uninduced and induced pATOM36 RNAi cell line were compared using a proteomic analysis (Fig. 3C, top panel). The experiment also showed that the levels of 11 OM proteins, including ATOM19 and ATOM46, were significantly reduced more than 1.5-fold in the induced RNAi cells. These proteins largely overlap with previously identified pATOM36 substrates (6).

**Figure 3.**
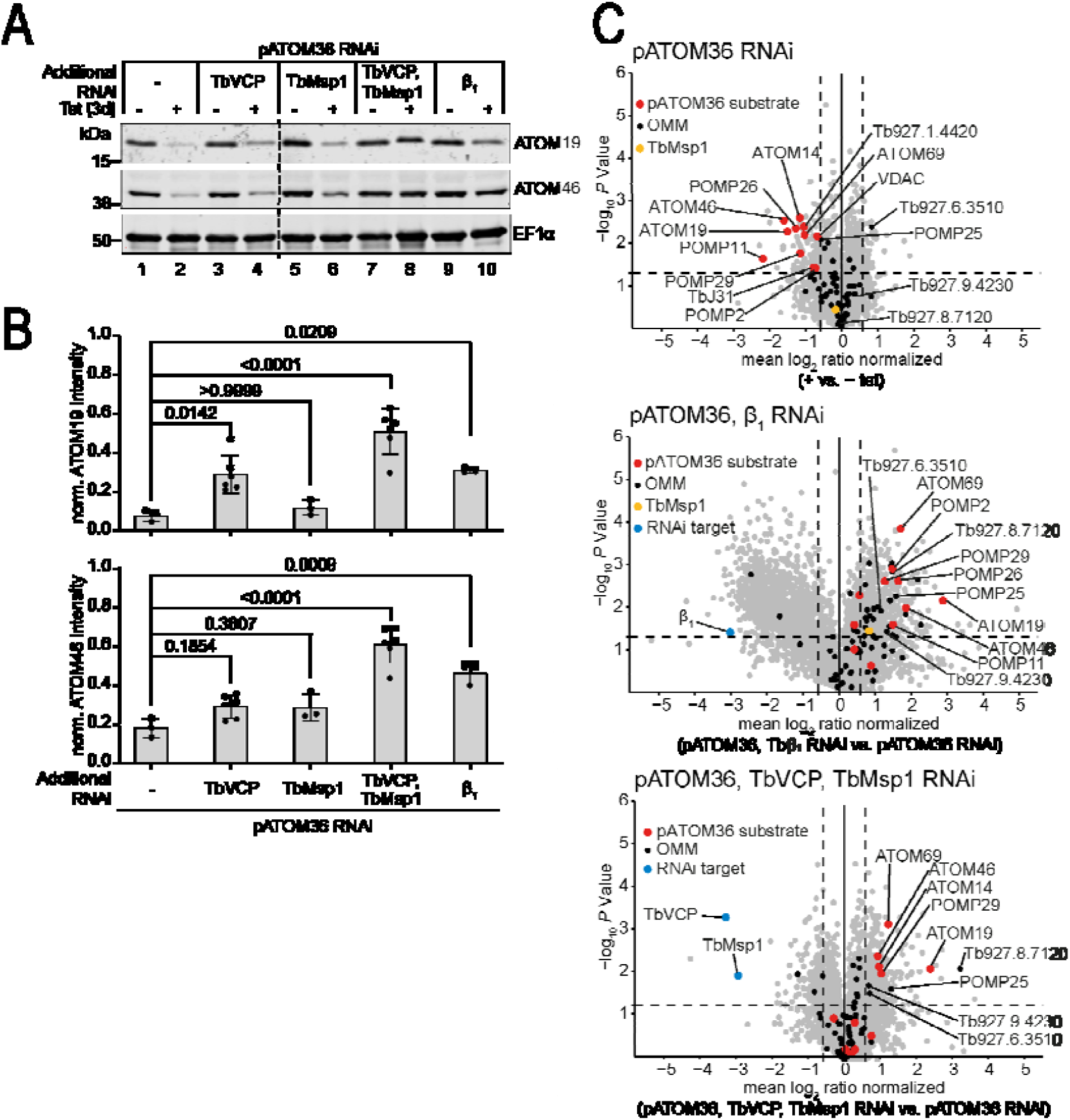
TbVCP and TbMsp1 are synergistically involved in the degradation of pATOM36 substrates by the cytosolic proteasome. A. Western blot analysis of total cellular extract (3×10^6^ cells each) of the indicated uninduced and induced single, double and triple RNAi cell lines (-/+ Tet), probed with ATOM19 and ATOM46 antisera. EF1α serves as loading control. B. Quantifications of ATOM46 and ATOM19 levels in the RNAi cell lines from immunoblots shown in A. The signal for each sample was _abeling_n to its respective EF1α signal and then to the respective signal in uninduced cells. Data are presented as mean values with error bars corresponding to the standard deviation (n=3 to 6). The p values indicated in the graph were calculated using a one-way ANOVA followed by a Bonferroni post hoc test to allow for multiple comparisons. C. Volcano plots _abeling_ng quantitative MS data of whole cell extracts from the indicated RNAi cell lines (n=3). Relative protein quantification was based on peptide stable isotope dimethyl _abeling. Shown are comparisons of uninduced and induced pATOM36 RNAi cells (top), induced pATOM36 RNAi cells and induced pATOM36/proteasome subunit β1 double RNAi cells (middle), and induced pATOM36 RNAi cells and induced pATOM36/TbVCP/TbMsp1 triple RNAi cells (bottom).

To test whether destabilised pATOM36 substrates are digested by the cytosolic proteasome, we produced a cell line able to knockdown both pATOM36 and the proteasomal subunit β1 (for a characterisation of all double and triple RNAi cell lines used in this study see Fig. S5). A comparison of this pATOM36/subunit β_1_ double RNAi cell line with the single pATOM36 RNAi cell line by immunoblot analysis indicated that the levels of ATOM46 and ATOM19 were significantly stabilised (Fig. 3A, compare lanes 2 and 10). Three-to-five-fold more ATOM19 and ATOM46 was found in cells depleted for pATOM36 and proteasomal subunit β_1_ in comparison to cells only depleted for pATOM36. This was in line with data from a quantitative proteomics analysis of induced samples of the same two cell lines (Fig. 3C, middle panel), which showed a significant more than 1.5-fold enrichment of seven pATOM36 substrates, including ATOM19 and ATOM46, indicating that their levels were stabilised. Moreover, a number of other OM proteins not previously shown to be substrates of pATOM36 were also stabilised. We conclude from this experiment that pATOM36 depletion triggers a pathway which feeds destabilised pATOM36 substrates to the cytosolic proteasome.

### TbMsp1 and TbVCP are implicated in proteasomal degradation of pATOM36 substrates

How can the cytosolic proteasome access membrane-integral pATOM36 substrates? In Opisthokonts, the AAA-ATPase Msp1 is able to extract TA proteins from the OM (46). Thus, we decided to test whether TbMsp1 could be involved in the degradation of the integral OM proteins ATOM19 and ATOM46 in pATOM36-depleted cells using the same approach that was used to show the involvement of the proteasome. However, in contrast to the pATOM36/subunit β_1_ double RNAi cell line (Fig. 3A, compare lanes 9 and 10), combining TbMsp1 RNAi with pATOM36 RNAi (Fig. 3A, compare lanes 5 and 6) did not significantly prevent the degradation of ATOM19 and ATOM46.

In Opisthokonts, the AAA-ATPase VCP is involved in various pathways that remove OM proteins from their membrane to allow for their degradation (46). In order to find out whether TbVCP, the trypanosomal VCP homolog (47, 48), plays a similar role in the pATOM36 triggered pathway, we produced a double RNAi cell line allowing simultaneous depletion of pATOM36 and TbVCP (Fig. S5). Immunoblot analyses of this cell line showed that, while the level of ATOM19 was slightly yet significantly stabilised upon pATOM36 and TbVCP depletion in comparison to the level found in pATOM36-depleted cells, the same was not the case for ATOM46 (Fig. 3A, B). Thus, simultaneous ablation of pATOM36 and TbVCP gave essentially the same results that were observed in the pATOM36/TbMsp1 double RNAi cell line.

These results can best be explained if the depletion of one AAA-ATPase protein, TbMsp1 or TbVCP, allowed its activity to be at least partially compensated by the other. To directly test this hypothesis, we generated a triple RNAi cell line, targeting pATOM36, TbMsp1 and TbVCP simultaneously (Fig. S5). With this cell line we could show that depletion of all three proteins significantly restored the levels of the ATOM19 and ATOM46 to approximately 3-6 fold of their levels in pATOM36-depleted cells (Fig. 3A, compare lanes 7 and 8). These results were independently confirmed and extended by a complementary proteomic analysis which compared the induced pATOM36 cell line (corresponding to lane 2 in Fig. 3A) with the induced triple RNAi cell line depleting pATOM36, TbMsp1 and TbVCP1 simultaneously (corresponding to lane 8 in Fig. 3A). In this experiment, five pATOM36 substrates and a few other OM proteins were significantly enriched more than 1.5-fold indicating that their levels were stabilised (Fig. 3C, bottom panel). The simplest explanation for these results is that TbMsp1 and TbVCP have redundant, at least partially, synergistic functions in the MAD pathway that leads to the degradation of pATOM36-dependent substrates.

### TbMsp1 interactors contribute to the function of the MAD pathway

Using the same approach, it was possible to test whether the four mitochondrial OM proteins that we identified to be in the same protein complex as Msp1 played a functional role in the MAD pathway investigated in this study. We constructed a series of triple RNAi cell lines, depleting either POMP31, POMP19, TbJ31 or TbTsc13 together with pATOM36 and TbVCP, to trigger the MAD pathway and to prevent pATOM36 substrates being degraded via TbVCP-mediated extraction from the OM (Fig. S5). Upon induction of RNAi, a significant restoration in the levels of ATOM19 and ATOM46 was detectable by immunoblot in triple RNAi cell lines where either TbJ31, POMP31 and TbTsc13 were depleted along with pATOM36 and TbVCP, in comparison to cells in which only pATOM36 and TbVCP were depleted (Fig. 4A, compare lanes 2 with lanes 6, 8, 10, Fig. 4B). In the case of TbJ31, only the level of ATOM46 restoration was significant. This observed restoration in the levels of ATOM19 and ATOM46 upon pATOM36, TbVCP and either TbJ31, POMP31 or TbTsc13 depletion phenocopies the effects observed in the triple RNAi cell line targeting pATOM36, TbVCP1 and TbMsp1. This strongly suggests that TbJ31, POMP31 and TbTsc13 do not only form a stable complex with mitochondrial TbMsp1, but that each of the three proteins also contributes to TbMsp1 function in the MAD pathway triggered by pATOM36 depletion. The triple RNAi cell line depleted for POMP19 did not significantly restore the levels of ATOM46 or ATOM19, suggesting that it does not affect mitochondrial TbMsp1 activity.

**Figure 4.**
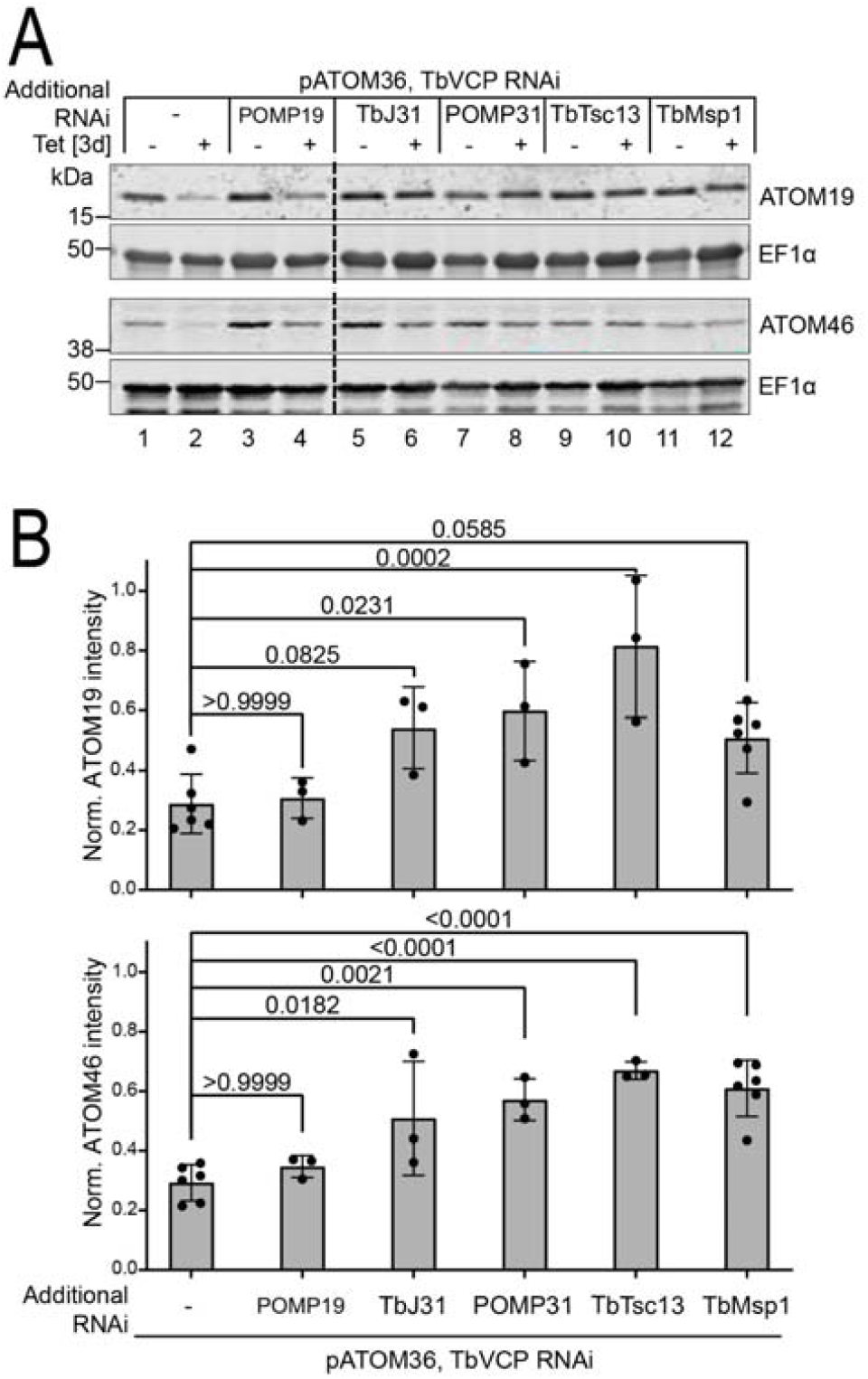
TbJ31, POMP31 and TbTsc13 are required for the degradation of pATOM36 substrates by TbMsp1. A) Western blot analysis of total cellular extracts (3×10^6^ cells each) of the indicated uninduced and induced double and triple RNAi cell lines (-/+ Tet), probed with ATOM19 and ATOM46 antisera. EF1α was used as a loading control. B)Quantifications of ATOM46 and ATOM19 levels in the RNAi cell lines from immunoblots shown in Fig. 3A. The signal for each sample was normalised to its respective EF1α signal and then to the respective level in uninduced cells. Data are presented as mean values with error bars corresponding to the standard deviation (n=3-6). The p values indicated in the graph were calculated using a one-way ANOVA followed by a Bonferroni post hoc test to allow for multiple comparisons.

## Discussion

We have discovered a pathway in *T. brucei* that removes destabilised α-helically anchored proteins from the mitochondrial OM. This pathway is triggered upon depletion of the OM protein biogenesis factor pATOM36. Previous studies suggest that pATOM36 has two distinct functions. It mediates the integration of ATOM46 and ATOM19 into the heterooligomeric ATOM complex after the proteins have been inserted into the OM (6). However, it can also facilitate insertion of certain proteins into the OM, as was shown for POMP10 (5). Removal of pATOM36 prevents integration of several ATOM subunits into the ATOM complex, leading to their degradation by the cytosolic proteasome. This degradation requires their selective extraction from the OM by the trypanosomal homologs of the Opisthokont mitochondrial quality control components Msp1 or VCP. There is some redundancy in the system as knocking down only one of the two AAA-ATPases hardly affects the pathway. Thus, this newly discovered TbMsp1 and TbVCP-linked pathway allows trypanosomes to safeguard their OM functions, maintaining this essential interface for mitochondrial-intracellular communications.

Maintenance of protein homeostasis is essential to maintain cellular functions under both unstressed and stress conditions. Many membrane proteins in eukaryotes require selective trafficking to specific subcellular compartments and assembly into defined stoichiometric complexes before functioning. This assembly process is not 100% efficient and thus degradation of unassembled, potentially harmful complex subunits is required. Msp1 is known to extract these orphaned proteins from both the mitochondrial OM as well as the peroxisomal membrane, allowing their degradation by the proteasome (15, 18, 49). In yeast, Msp1 has been shown to be sufficient for membrane protein extraction (5), or, in case of particular substrates, to function together with an interacting protein that is induced by a specific trigger, for example in MitoCPR (19).

Trypanosomal TbMsp1 surprisingly forms a stable complex with at least four other integral mitochondrial OM proteins. Three of these interactors, POMP31, TbJ31 and TbTsc13 contribute to TbMsp1 activity in the MAD pathway that is triggered upon pATOM36 depletion. While POMP31 is only found in kinetoplastids, TbJ31 is an orthologue of the mammalian mitochondrial OM J-protein, DNAJC11, although it lacks a complete HPD motif and thus is a J-like protein (43). TbJ31 and DNAJC11 both have a C-terminal DUF3395 domain suggested to mediate protein-protein interactions (50). It has also been reported that mammalian DNAJC11 may transiently interact with the mitochondrial contact site and cristae organizing system (MICOS) complex (50, 51). TbTsc13 shows similarity to the enoyl-CoA reductase of the ER elongase complex (52–55). Interestingly, it has an N-terminal ubiquitin-like domain that is exposed to the cytosol (56). As yet, we do not understand the specific role these TbMsp1-interacting proteins may play in the described MAD pathway. However, the notion that TbMsp1 may act in concert with a J-like protein that could directly or indirectly regulate chaperones seems plausible in this context. The same is the case for the ubiquitin-like domain of TbTsc13, which potentially could facilitate proteasome binding and activation (57).

How pATOM36 substrates are recognised by the MAD pathway is not yet understood, in particular we do not know how these proteins can be recognised efficiently by both TbMsp1 and TbVCP. Msp1 substrate specificity is known to be multifaceted (58), however as the pATOM36 substrates we focused on in this work, ATOM46 and ATOM19, are integral parts of the ATOM complex, we could hypothesise that these proteins become orphaned upon pATOM36 depletion allowing them to become substrates of TbMsp1. Nevertheless, not all pATOM36 substrates are known to be components of multiprotein complexes. Cytosolic VCP is involved in diverse cellular processes, and its substrate specificity in other organisms is governed by its numerous cofactors, many of which interact with ubiquitin conjugated to its substrates (59, 60). The potential requirement for selective ubiquitination cascades adds another layer of yet undefined diversity to the regulation of this process.

Understanding variations in mitochondrial biogenesis across eukaryotes can provide insight into their evolution as well as into the process of how the endosymbiotic bacterial ancestor of the mitochondrion converted into an organelle. VCP and Msp1 are conserved throughout eukaryotes, and thus were present in the last eukaryotic common ancestor (LECA). However, the convergent evolution of known divergent OM protein biogenesis factors (pATOM36, MTCH2 and MIM) for α-helically anchored OM proteins between, and even within, distinct eukaryotic supergroups suggests that LECA did not contain a protein with this function (7).

This is in agreement with the notion that LECA contained a much simpler β-barrel-based OM protein import system (61, 62), whereas most additional α-helical subunits of the TOM complex, for example the receptors, were added later after a first divergence of eukaryotes to confer specificity and efficiency of the import process (61, 63–65). Thus, the role of Msp1 in removing orphan α-helical OM proteins is likely not its ancestral one. Instead, the requirement of Msp1 to clear precursor blockages in the OM protein import machinery may have evolved first (19). Whether TbMsp1 has retained this activity remains to be investigated.

Thus, the TbMsp1 function linked to surveillance of OM protein biogenesis likely arose after pATOM36 evolution, and the same is the case for the mitochondrial OM protein complex formed by Msp1 and its interactors, three of which contribute to its activity. The MAD pathway triggered by the depletion of pATOM36 is to our knowledge the first one to be characterised in any eukaryote that is specifically linked to defects in the OM protein biogenesis.

If the emergence of pATOM36 drove the evolution of a Msp1/VCP-linked pathway to survey and maintain the integrity of its activity, did the same happen in Opisthokonts? Intriguingly, there are hints that depletion of yeast MIM or mammalian MTCH2 may drive MAD pathways. Loss of these proteins does result in depletion in the level of at least some of their substrates (7, 8), reminiscent of the proteasomal degradation of pATOM36 substrates by the MAD pathway described here. Accumulation of orphan OM proteins is likely harmful for all mitochondria, suggesting that a pathway to deal with such proteins might be required in all eukaryotes.

We therefore expect that the independent establishment of specific OM protein biogenesis pathways in different phylogenetic groups resulted in the parallel evolution of the corresponding MAD pathways in the same groups. It is likely that these systems are also connected to the widely conserved AAA-ATPases Msp1 and VCP. Should this be the case, it will be interesting to find out whether they, as with TbMsp1, also require additional factors for full activity and, if yes, what their identity might be.

There has been much progress in defining mitochondrial quality control pathways in Opisthokonts such as yeast and metazoans. However, only very recently have studies on mitochondrial quality control expanded out of this narrow range of eukaryotic diversity. A MAD pathway has been found in trypanosomes, a member of the Discoba supergroup, that facilitates the removal of mistargeted aggregation-prone mitochondrial proteins from the cytosol (66). The results suggested that the depletion of cytosolic chaperones may be a general trigger of MAD throughout eukaryotes. The present Msp1 and VCP-linked pathway is the second MAD pathway discovered in trypanosomes. Further studies in other non-classical model systems are expected to improve our understanding of the fundamental features of such pathways, which are similar not due to common descent, but because all eukaryotes have to cope with the shared constraints imposed by hosting mitochondria.

## Materials and Methods

### Reagents and Tools Table

| **Reagent type (species) or resource** | **Designation** | **Source or reference** | **Identifiers** | **Additional information** |
| --- | --- | --- | --- | --- |
| Cell line (T.brucei) | 29.13, procyclic | (67) |  | Wild type (WT) |
| Antibody | Anti-HA (mouse, monoclonal) | BioLegend | 901503 (MMS-101R) | WB (1:5000), IFA (1:1000) |
| Antibody | Anti-myc (mouse, monoclonal) | Invitrogen | 132500 | WB (1:2000), IFA (1:50) |
| Antibody | Anti-ATOM19 | Eurogentec | Polyclonal antibody against purified protein | WB (1:500) |
| Antibody | Anti-ATOM46 | (64) |  | WB (1:500) |
| Antibody | Anti-pATOM36 | (7) |  | WB (1:250) |
| Antibody | Anti-TbVCP | Gift from Prof. James Bangs, SUNY Buffalo, US |  | WB (1:50) |
| Antibody | Anti-ALD | Gift from Paul Michels, University of Edinburgh, Scotland |  | IFA (1:1500) |
| Antibody | Anti-BiP | Gift from Prof. James Bangs, SUNY Buffalo, US |  | IFA (1:2500) |
| Antibody | Anti-TbMsp1 | Eurogentec | Peptide Antibody against C+DEALKRVRPSMASSV | WB (1:1000) |
| Antibody | Anti-ATOM40 (rabbit, polyclonal) | (33) |  | Bleed 1, IFA (1:1000) |
| Antibody | Anti-VDAC (rabbit, polyclonal) | (33) |  | WB (1:1000) |
| Antibody | Anti-EF1α<br>(mouse,<br>monoclonal) | Merk Millipore | 05-235 | WB (1:10000) |
| Antibody | Anti-cytochrome<br>c (rabbit,<br>polyclonal) | (68) |  | WB (1:100) |
| Antibody | Anti-mouse<br>IRDye 680LT<br>conjugated<br>(goat) | LI-COR<br>Biosciences | PN 926-68020 | WB (1:20000) |
| Antibody | Anti-rabbit<br>IRDye 800CW<br>conjugated<br>(goat) | LI-COR<br>Biosciences | PN 926-32211 | WB (1:20000) |
| Antibody | Goat Anti-mouse Alexa<br>Fluor 596 | ThermoFisher<br>Scientific | # A-11032 | IFA (1:1000) |
| Antibody | Goat Anti-rabbit<br>Alexa Fluor 488 | ThermoFisher<br>Scientific | # A-11008 | IFA (1:1000) |
| Commercial<br>assay or kit | Prime-a-Gene<br>labelling kit | Promega | U1100 | Radioactive labelling<br>of Northern probes |
| Commercial<br>assay or kit | EZView Red<br>Anti-c-myc<br>affinity gel | Sigma Aldrich | E6654 | CoIP |
| Commercial<br>assay or kit | Anti-HA affinity<br>matrix | Roche | 11815016001 | CoIP |
| Chemical<br>compound,<br>drug | Tetracycline<br>Hydrochloride | Sigma Aldrich | T7660 | Tet |
| Chemical<br>compound,<br>drug | Digitonin | Biosynth | 103203 | Generation of crude<br>mitochondrial<br>fractions |
| Chemical<br>compound,<br>drug | Paraformaldehy<br>de (PFA) | Fluka | UN2213 |  |
| Chemical<br>compound,<br>drug | Albumin (BSA)<br>Fraktion V (pH<br>7,0) | Applichem | A1391 |  |
| Chemical<br>compound,<br>drug | Triton® X-100 | Merck<br>Millipore | 108603 |  |
| Chemical<br>compound,<br>drug | Tween 20 | AppliChem | A4974 |  |
| Chemical<br>compound,<br>drug | Lysine-L U-13C,<br>U-15N (Lys8) | Euroisotop | CNLM-291-H | SILAC labelling |
| Chemical<br>compound,<br>drug | Arginine-L U-13C6, U-15N4<br>(Arg10) | Euroisotop | CNLM-539-H | SILAC labelling |
| Chemical compound, drug | Lysine-L, 4.4.5.5-D4 (Lys4) | Euroisotop | DLM-2640 | SILAC labelling |
| Chemical compound, drug | Arginine-L 13C6 (Arg6) | Euroisotop | CLM-2265-H | SILAC labelling |
| Chemical compound, drug | Formaldehyde, light (CH<sub>2</sub>O) | Sigma Aldrich | 252549 | Peptide stable isotope dimehtyl labelling |
| Chemical compound, drug | Formaldehyde, heavy (<sup>13</sup>CD<sub>2</sub>O) | Sigma Aldrich | 492620 | Peptide stable isotope dimehtyl labelling |
| Chemical compound, drug | Sodium cyano-borohydride (NaBH<sub>3</sub>CN) | Sigma Aldrich | 156159 | Peptide stable isotope dimehtyl labelling |
| Enzyme | Trypsin, MS approved | SERVA | 37286 |  |
| Software, algorithm | GraphPad Prism, version 6.0 f | Graphpad software | www.graphpad.com | Depiction of growth curves and analysis of western blot quantification |
| Software, algorithm | Fiji | ImageJ | doi:10.1038/nmeth.2019 | Processing of images |
| Software, algorithm | FigureJ | ImageJ Plugin | doi.org/10.1111/jmi.12069 | Assembly of microscopy figures |
| Software, algorithm | Image Studio Lite v. 5.2.5. | LI-COR Biosciences |  | Quantification of western blots |
| Software, algorithm | Adobe Illustrator | Adobe | www.adobe.com | Figure assembly |
| Software, algorithm | RStudio | RStudio | www.rstudio.com | Mass spectrometry analysis, volcano plotting |

#### Methods and Protocols Transgenic cell lines

Transgenic *T. brucei* cell lines were generated using the procyclic strain 29.13 (67). Cells were cultivated at 27°C in SDM-79 (69) supplemented with 10% (v/v) fetal calf serum (FCS), containing G418 (15 µg/ml, Gibco), hygromycin (25 µg/ml, InvivoGen), puromycin (2 µg/ml, InvivoGen), blasticidin (10 µg/ml, InvivoGen) and phleomycin (2.5 µg/ml, LifeSpan BioSciences) as required. RNAi or protein overexpression was induced by adding 1 µg/ml tetracycline to the medium.

To produce plasmids for ectopic expression of C-terminal triple c-myc-or HA-tagged TbMsp1 (Tb927.5.960), POMP31 (Tb927.6.3680), TbJ31 (Tb927.7.990), POMP19 (Tb927.10.510) and TbTsc13 (Tb927.3.1840), the complete ORFs of the respective gene were amplified by PCR and inserted in a modified pLew100 vector (67, 70) containing either a C-terminal triple c-myc-or HA-tag (71). One TbMsp1 allele was tagged in situ at the C-terminus with a triple HA-tag via a PCR approach, using a pMOTag vector containing a phleomycin resistance cassette as described in (71).

RNAi cell lines were prepared using a pLew100-derived vector with a 500bp target gene fragment and its reverse complement present with a 460 bp stuffer in-between, generating a stem-loop construct. The RNAis targeted the indicated nucleotides (nt) of the ORF of proteasome subunit β_1_ (nt 266-759), TbVCP (nt 423-896), TbTsc13 (nt 379-891), TbJ31 (nt 831-1255), POMP31 (nt 170-568), POMP19 (nt 182-610), TbMsp1 (nt 530-940). The pATOM36 RNAi construct was previously published (72).

#### Digitonin extraction

Cell lines were induced for 16 hr prior to the experiment to express the epitope-tagged proteins. Crude mitochondria-enriched fractions were obtained by incubating 1 ×10^8^ cells on ice in 0.6 M sorbitol, 20 mM Tris-HCl pH 7.5, 2 mM EDTA pH 8 containing 0.015% (w/v) digitonin for the selective solubilization of plasma membranes. Centrifugation (5 min, 6800 g, 4°C) yielded a cytosolic supernatant and a mitochondria-enriched pellet. Equivalents of 1.3 ×10^6^ cells of each fraction were analysed by SDS-PAGE and subsequent Western blotting to demonstrate organellar enrichment for proteins of interest.

#### Alkaline carbonate extraction

To separate soluble or peripherally membrane-associated proteins from integral membrane proteins, a mitochondria-enriched pellet was generated as described above by digitonin extraction and resuspended in 100 mM Na_2_CO_3_ pH 11.5. Centrifugation (10 min, 100000 g, 4°C) yielded a supernatant containing soluble proteins and a pellet containing membrane fragments. Equivalents of 7.5 ×10^6^ cells of each fraction were subjected to SDS-PAGE and immunoblotting.

#### Immunoprecipitation

Digitonin-extracted mitochondria-enriched fractions of 1×10^8^ induced cells were solubilized on ice in 20 mM Tris-HCl pH7.4, 0.1 mM EDTA, 100 mM NaCl, 25mM KCl, 1x protease inhibitor mix (Roche, EDTA-free) and 1% (w/v) digitonin. After centrifugation (15 min, 20817g, 4°C), the lysate (IN, input) was transferred to either 50 µl of HA bead slurry (anti-HA affinity matrix, Roche) or 50 µl c-myc bead slurry (EZview red anti-c-myc affinity gel, Sigma), both of which had been equilibrated in wash buffer (20 mM Tris-HCl pH 7.4, 0.1 mM EDTA, 100 mM NaCl, 10% glycerol, 0.2% (w/v) digitonin). After incubating at 4°C for 2 hr on a rotating wheel, the supernatant containing the unbound proteins (FT, flow through) was removed. The bead slurry was washed three times with wash buffer. Bound proteins were eluted by boiling the resin in 60 mM Tris-HCl pH 6.8 containing 2% SDS (IP). 2.5% of crude mitochondrial fractions (Input, IN) and unbound proteins in the flow through (FT), and 50% of the final eluates (IP) were separated by SDS-PAGE and analysed by western blot.

#### SILAC immunoprecipitations

Cells were grown for five days in SILAC medium (SDM80 containing 5.55 mM glucose, supplemented with 10% dialyzed, heat-inactivated FCS, 7.5 mg/l hemin) containing isotopically distinct variants of arginine (^12^C_6_ ^14^N_4_ /Arg0, ^13^C_6_ ^14^N_4_ /Arg6, or ^13^C_6_ ^15^N_4_ /Arg10; 226 mg/l each) and lysine (^12^C_6_ ^14^N_4_ /Lys0, ^12^C_6_ ^14^N_2_ ^2^H_4_ /Lys4, or ^13^C_6_ ^15^N_4_ /Lys8; 73 mg/l each) (Eurisotope). 2×10^8^ wildtype cells and cells expressing in situ tagged Msp1-HA were mixed and washed with 1x PBS. Crude mitochondria-enriched fractions were obtained by digitonin extraction as described above. The pellet of the digitonin extraction was subjected to immunoprecipitation as described above. Proteins were precipitated following the methanol-chloroform protocol (73) and further processed for liquid chromatography-mass spectrometry (LC-MS) analysis including reduction of cysteine residues, alkylation of thiol groups, and tryptic digestion as described before (74). The experiment was performed in three biological replicates with different labelling schemes.

#### RNA extraction and northern blotting

Acid guanidinium thiocyanate-phenol-chloroform extraction according to Chomczynski and Sacchi (75) was used for isolation of total RNA from uninduced and induced RNAi cells. Total cellular RNA was separated on a 1% agarose gel in 20 mM MOPS buffer supplemented with 0.5% formaldehyde. Northern probes were generated from gel-purified PCR products corresponding to the RNAi inserts and radioactively labelled using the Prime-a-Gene labelling system (Promega).

#### Immunofluorescence microscopy

Induced ×10^6^ cells overexpressing the indicated tagged proteins were harvested by centrifugation (5 min, 1800 g) and washed with 1x PBS. After resuspension in 1x PBS, the cells were left adhering on a glass slide in a wet chamber. The cells were fixed with 4% PFA, permeabilised with 0.2% Triton X-100 and blocked with 2% BSA in 1x PBS. Antibodies were incubated on the slides in 1% BSA, 1x PBS. The dried slides were mounted with Vectashield (containing 4, 6-diamidino-2-phenylindole (DAPI) (Vector Laboratories, P/N H-1200). Images were acquired with a DFC360 FX monochrome camera (Leica Microsystems) mounted on a DMI6000B microscope (Leica Microsystems). Image analysis was done using LASX software (version 3.6.20104.0, Leica Microsystems). The acquired images were processed using Fiji (ImageJ version 2.10./1.53; Java 1.8.0_172 [64 bit]) and composed to a Figure using FigureJ (76).

#### Peptide stable isotope dimethyl labelling and high-pH reversed-phase fractionation

RNAi cell lines were grown in triplicate in SDM-79 for three days, in the presence or absence of tetracycline. 1 ×10^8^ cells were centrifuged (8 min, 1258 g, RT) and washed with 1x PBS. The pellets were flash frozen in liquid nitrogen and subsequently processed for tryptic in-solution digestion as described before (77). Dried peptides were reconstituted in 100 mM tetraethylammonium bicarbonate, followed by differential labelling with “light” or “heavy” formaldehyde (CH_2_ O/_13_CD_2_ O; Sigma-Aldrich) and sodium cyanoborohydride (NaBH_3_CN; Sigma-Aldrich) (78). Labelling efficiencies (> 99% for all individual experiments) were determined by LC-MS analysis. Equal amounts of differentially “light” and “heavy” labelled peptides derived from the respective control and induced RNAi cells were mixed, purified and fractionated by high pH reversed-phase chromatography using StageTips essentially as described previously (79). In brief, peptides, reconstituted in 10 mM NH_4_OH, were loaded onto StageTips and eluted stepwise with 0%, 2.7%, 5.4%, 9.0%, 11.7%, 14.4%, 36% and 65% (v/v each) acetonitrile (ACN)/10 mM NH_4_OH. Fractions 1 and 7 (0% and 36% ACN eluates) and fractions 2 and 8 (2.7% and 65% ACN eluates) were combined for LC-MS analysis.

#### Quantitative LC-MS analysis

Prior to LC-MS analysis, peptides were desalted using StateTips, vacuum-dried, and reconstituted in 0.1% (v/v) trifluoroacetic acid. LC-MS analyses were performed using either a Q Exactive Plus (Msp1-HA SILAC IPs) or an Orbitrap Elite (RNAi experiments) mass spectrometer connected to an UltiMate 3000 RSLCnano HPLC system (all instruments from Thermo Fisher Scientific, Germany). Peptides were loaded and concentrated on PepMap C18 precolumns (length, 5 mm; inner diameter, 0.3 mm; Thermo Scientific) at a flow rate of 30 µl/min and separated using Acclaim PepMap C18 reversed-phase nano-LC columns (length, 500 mm; inner diameter, 75 μm; particle size, 2 μm; pore size, 100 Å; Thermo Scientific) at a flow rate of 0.25 µl/min. The solvent system used for the elution of peptides from Msp1-HA SILAC IP experiments consisted of 0.1% (v/v) formic acid (FA; solvent A1) and 86% (v/v) ACN/0.1% (v/v) FA (solvent B1). The following gradient was applied: 4% - 39% solvent B1 in 195 min followed by 39% - 54% B1 in 15 min, 54% - 95% B1 in 3 min, and 5 min at 95% B1. For the elution of peptides from RNAi experiments, 4% (v/v) dimethyl sulfoxide (DMSO)/0.1% (v/v) FA (solvent A2) and 48% (v/v) methanol/30% (v/v) ACN/4% (v/v) DMSO/0.1% (v/v) FA (solvent B2) were used. A gradient ranging from 3% - 65% solvent B2 in 65 min, 65% - 80% B2 in 5 min, and 5 min at 80% B2 was applied.

Mass spectrometric data were acquired in data-dependent mode. The Q Exactive Plus was operated with the following settings: mass range, *m/z* 375 to 1,700; resolution, 70,000 (at *m/z* 200); target value, 3× 10^6^; and maximum injection time (max. IT), 60 ms for MS survey scans. Fragmentation of up to 12 of the most intense multiply charged precursor ions by higher-energy collisional dissociation was performed with a normalized collision energy (NCE) of 28%, a target value of 10^5^, a max. IT of 120 ms, and a dynamic exclusion (DE) time of 45 s. The parameters for MS analyses at the Orbitrap Elite were as follows: mass range, *m/z* 370 to 1,700; resolution, 120,000 (at *m/z* 400); target value, 10^6^; and max. IT, 200 ms for survey scans. A TOP15 (pATOM36/subunit β_1_ double and pATOM36/TbVCP/TbMsp1 triple RNAi experiments) or TOP25 (pATOM36 RNAi experiments) method was applied for fragmentation of multiply charged precursor ions by low energy collision-induced dissociation in the linear ion trap (NCE, 35%; activation q, 0.25; activation time, 10 ms; target value, 5,000; max. IT, 150 ms; DE, 45 s). Proteins were identified and quantified using MaxQuant/Andromeda (80, 81) (version 1.5.5.1 for Msp1-HA SILAC IP and 1.6.0.1 for RNAi data). Mass spectrometric raw data were searched against a TriTryp database specific for *T. brucei* TREU927 (release version 8.1 for Msp1-HA SILAC IP and 36 for RNAi data; downloaded from https://tritrypdb.org). For protein identification, MaxQuant default settings were applied, with the exception that only one unique peptide was required. For relative quantification, the appropriate settings for SILAC labeling (light labels, Lys0/Arg0; medium-heavy, Arg6/Lys4; heavy, Lys8/Arg10) or stable isotope dimethyl labeling (light, dimethylLys0/dimethylNterLys0; heavy, dimethylLys6/dimethylNterLys6) were chosen. Quantification was based on at least one ratio count. The options ‘match between runs’ and ‘requantify’ were enabled. Only proteins quantified in at least two independent replicates per dataset were considered for further analysis. The mean log_10_ (SILAC IP data) or mean log_2_ (RNAi data) of protein abundance ratios was determined and a one-sided (SILAC IP data) or two-sided (RNAi data) Student’s t-test was performed. For information about the proteins identified and quantified, see PXD039631 (Msp1-HA SILAC IPs) and PXD039634 (RNAi experiments) in the PRIDE database.

#### Computational analysis of proteins

Conserved structural elements of Msp1 (82–84) are highlighted in Fig. 1A. Transmembrane domains were predicted using Phobius (85) (TbMsp1, POMP31, TbJ31, POMP19) or HMMTOP (86) (TbTsc13), conserved domains were either predicted with ncbi.nlm.nih.gov/Structure (POMP19, TbTsc13) or annotated Pfam domains on HMMER (87) (TbMsp1, TbJ31). The ubiquitin-like domain of TbTsc13 was predicted by HHpred (88). The multiple amino acid sequence alignment of TbMsp1, ATAD1 from *H. sapiens* (HsATAD1), and Msp1 from *S. cerevisiae* (ScMsp1) shown in Fig. S1 was performed with Clustal Omega (89).

#### Data availability

The mass spectrometry data have been deposited to the ProteomeXchange Consortium (90) via the PRIDE (91) partner repository and are accessible using the dataset identifiers PXD039631 (SILAC-IP data) and PXD039634 (RNAi data).

Msp1-HA SILAC IPs data:

Accession number: PXD039631

Username

Password: g3gWYtw1

RNAi data:

Accession number: PXD039634

Username

Password: CE0IUa0i

## Acknowledgements

We thank Bettina Knapp for technical assistance. We thank Noemis Zbären and Gabriel Kleese for help at the initial stage of the project. Work in the lab of B.W. was supported by the Deutsche Forschungsgemeinschaft (DFG, German Research Foundation) project ID 403222702/SFB 1381 and Germany’s Excellence Strategy (CIBSS – EXC-2189 – Project ID 390939984). Work in the lab of A.S. was supported in part by NCCR RNA & Disease, a National Centre of Competence in Research (grant number 205601) and by project grant SNF 205200 both funded by the Swiss National Science Foundation.

## Conflict of interest

The authors declare that they have no conflict of interest.

## Supplemental Figures

**Supplementary Figure 1.**
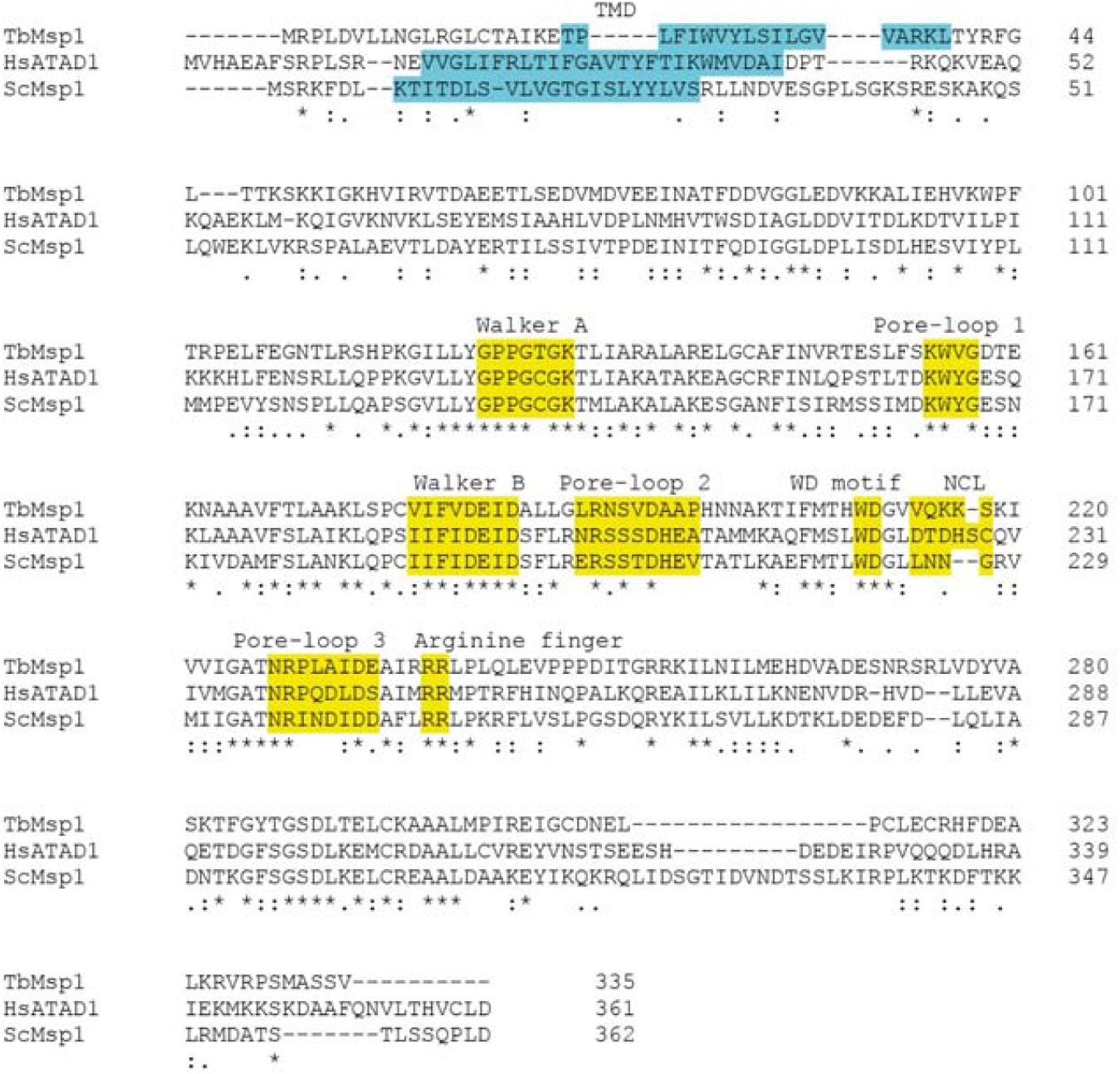
TbMsp1 contains motifs conserved across species. Clustal Omega multiple sequence alignment of amino acid sequences of TbMsp1, ATAD1 from H. sapiens (HsATAD1), and Msp1 from S. cerevisiae (ScMsp1). Predicted TMDs are coloured in blue. The conserved structural elements coloured in yellow are: (i) the Walker A and B motif for nucleotide binding and ATP hydrolysis, (ii) three pore-loop motifs, which are involved in gripping and unfolding substrates and driving translocation through the pore, (iii) a WD motif, which likely contributes to the coupling of ATP hydrolysis to conformational changes required for successful substrate translocation, (iv) a nucleotide communication loop (NCL) and (v) an arginine finger motif.

**Supplementary Figure 2.**
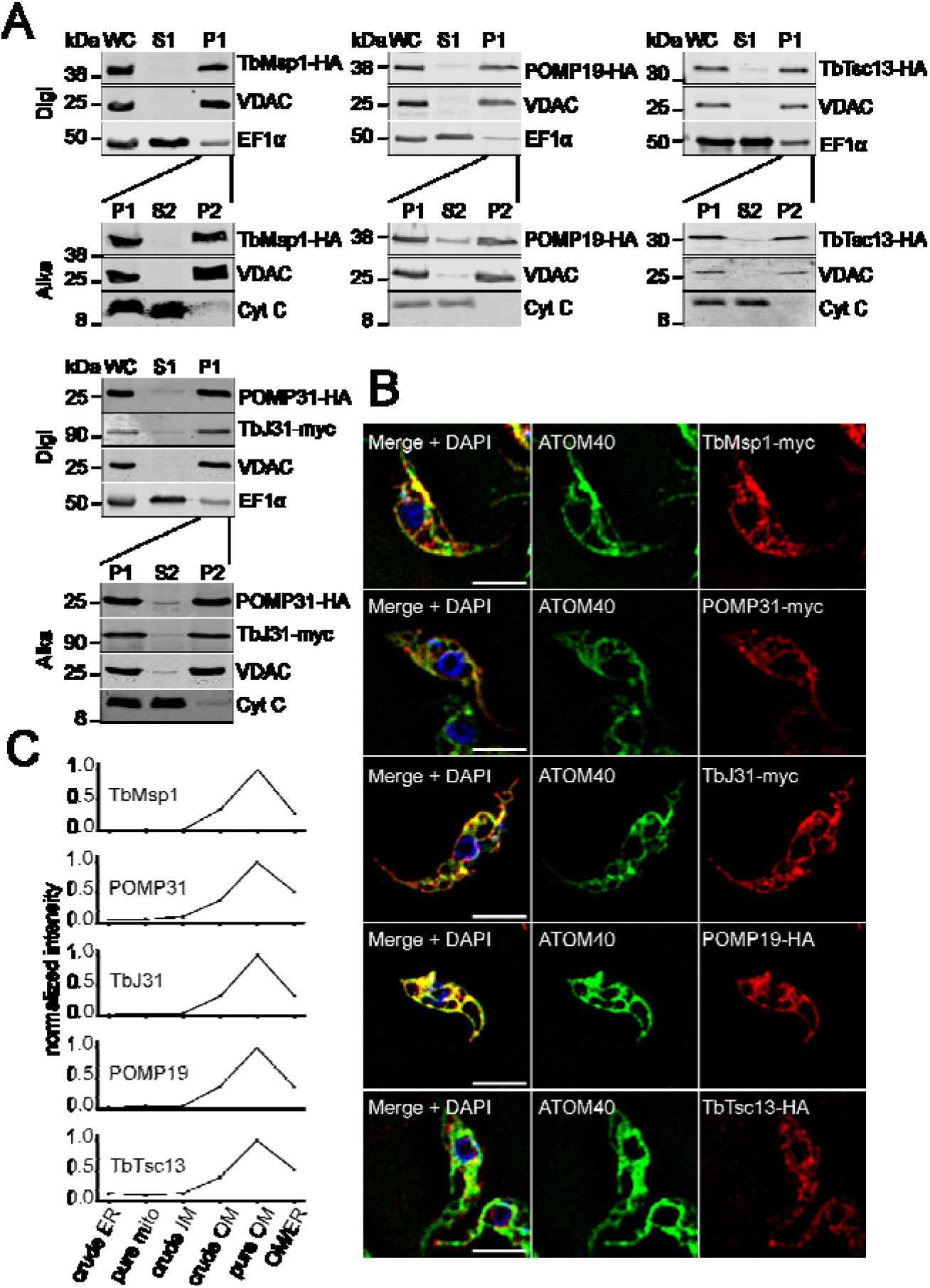
TbMsp1, POMP31, TbJ31, POMP19 and TbTsc13 are integral OM proteins. A. Immunoblot analysis of whole cells (WC), soluble cytosolic (S1) fractions and digitonin-extracted mitochondria-enriched (P1) from cells overexpressing the indicated C-terminally myc-or HA-tagged proteins. Immunoblots were probed with anti-tag antibodies and antisera against VDAC and EF1α, which serve as markers for mitochondria and cytosol, respectively. P1 fractions were subjected to alkaline carbonate extraction at pH 11.5 resulting in soluble supernatant (S2) and membrane-enriched pellet (P2) fractions. Immunoblots were probed with anti-tag antibodies and antisera against VDAC and cytochrome c (cyt _c_), which serve as markers for integral and peripheral membrane proteins, respectively. B.Immunofluorescence analysis of whole cells overexpressing the indicated C-terminally myc or HA-tagged proteins detected with anti-tag antibodies. ATOM40 serves as a mitochondrial marker. DAPI marks both nuclear and mitochondrial DNA in merged images. Scale bars: 5 μm. C.Normalized abundance profile of TbMsp1, POMP19, POMP31, TbJ31 and TbTsc13 over six subcellular fractions from a previously published proteomic analysis showing maximal intensity in the OM fraction (33).

**Supplementary Figure 3.**
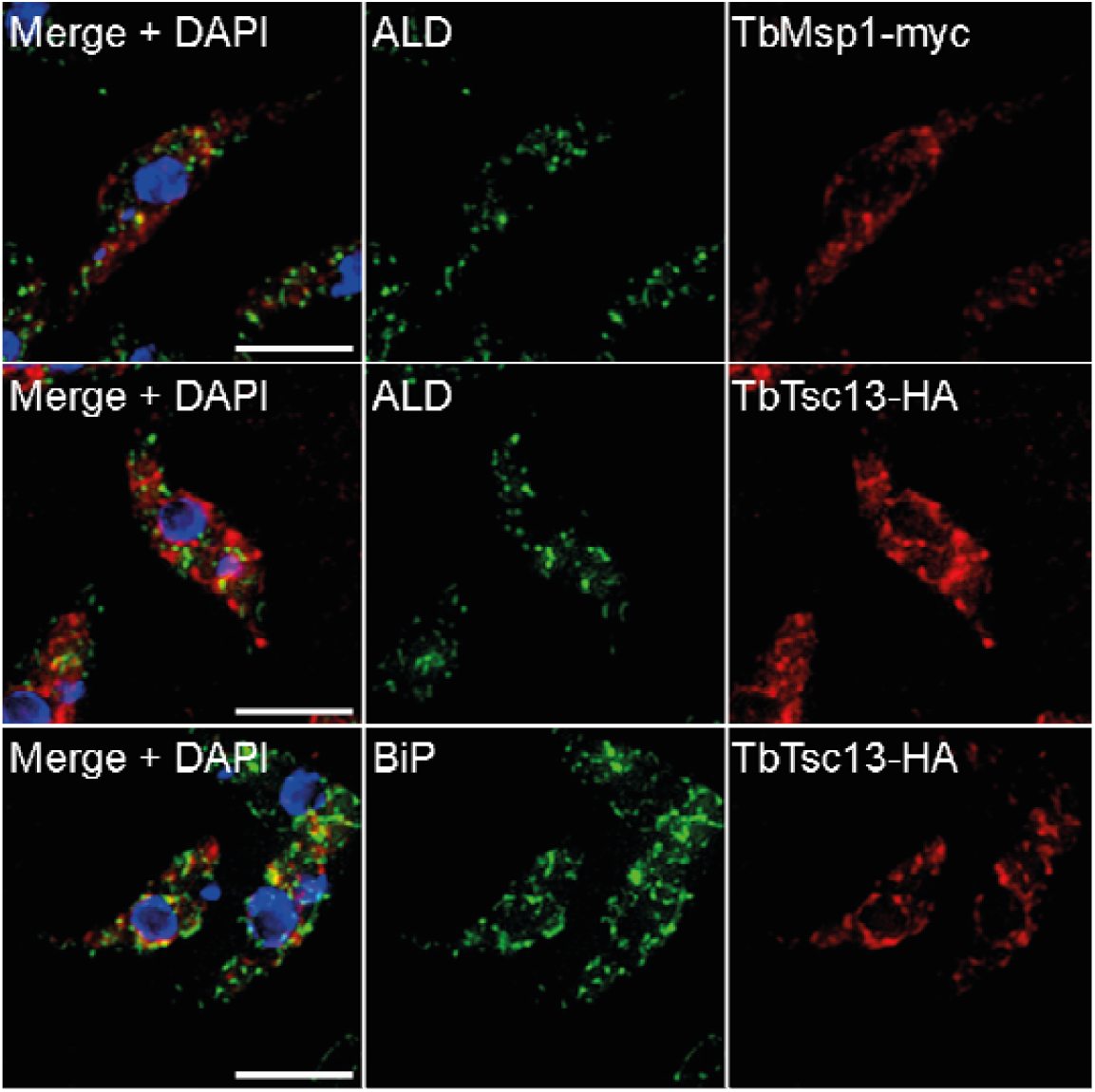
TbTsc13 and TbMsp1 are partially localized to the ER or glycosomes, respectively. Immunofluorescence analysis of whole cells overexpressing the indicated C-terminally myc or HA-tagged proteins detected with anti-tag antibodies. Staining for ALD and BiP serve as glycosomal and ER markers respectively. DAPI marks both nuclear and mitochondrial DNA in merged images. Scale bars: 10 μm.

**Supplementary Figure 4.**
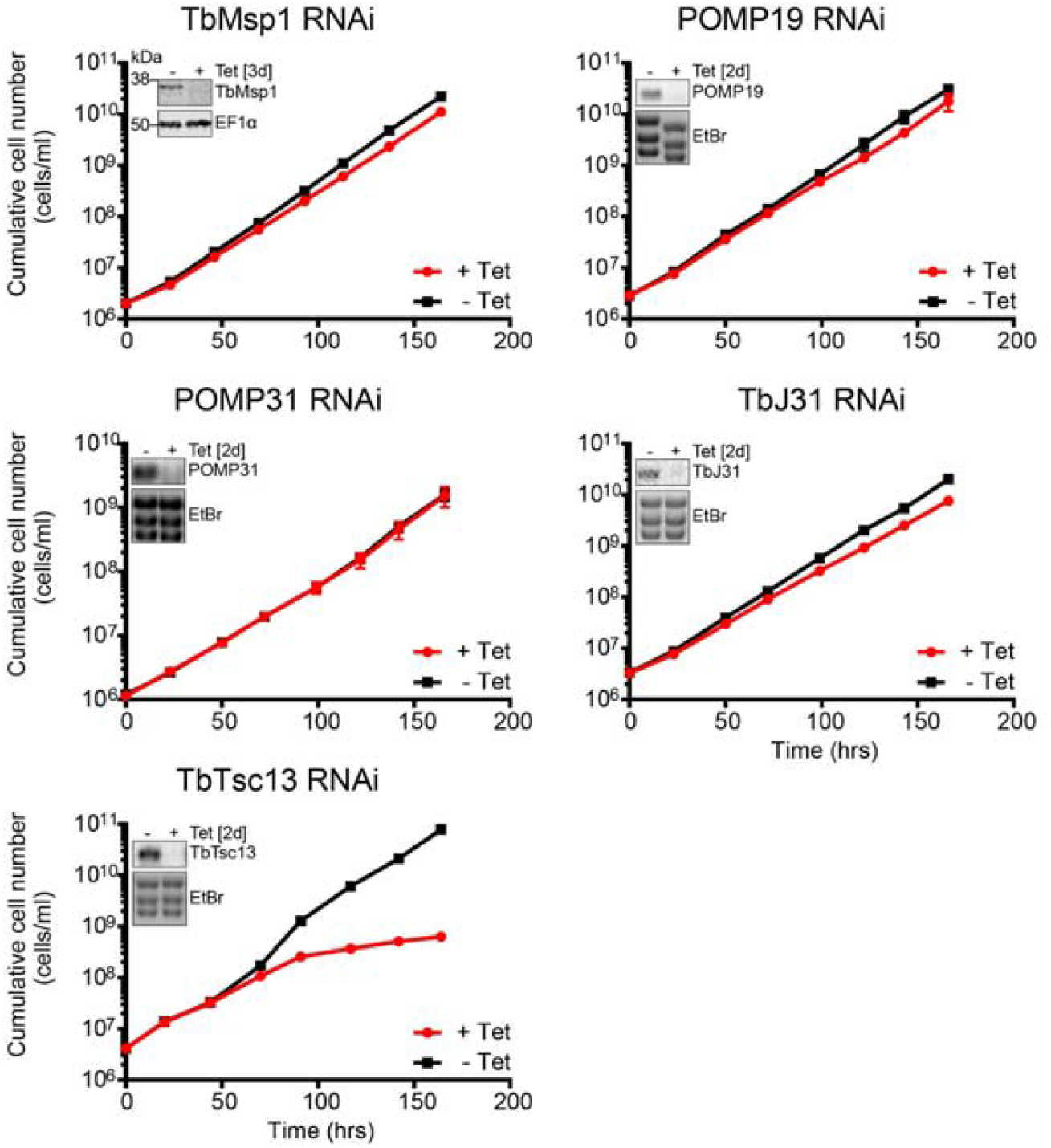
Verification and growth analysis of TbMsp1, POMP19, POMP31, TbJ31 and TbTsc13 RNAi cell lines. Growth curve of the indicated induced (+Tet) and uninduced (-Tet) RNAi cell lines. Error bars correspond to the standard deviation (n = 3). The inset panels show the efficiency of RNAi for the indicated cell lines, either three days after induction when analysed by western blot or two days after induction when analysed by northern blot. EF1α or ethidium bromide stained rRNAs serve as loading controls respectively.

**Supplementary Figure 5.**
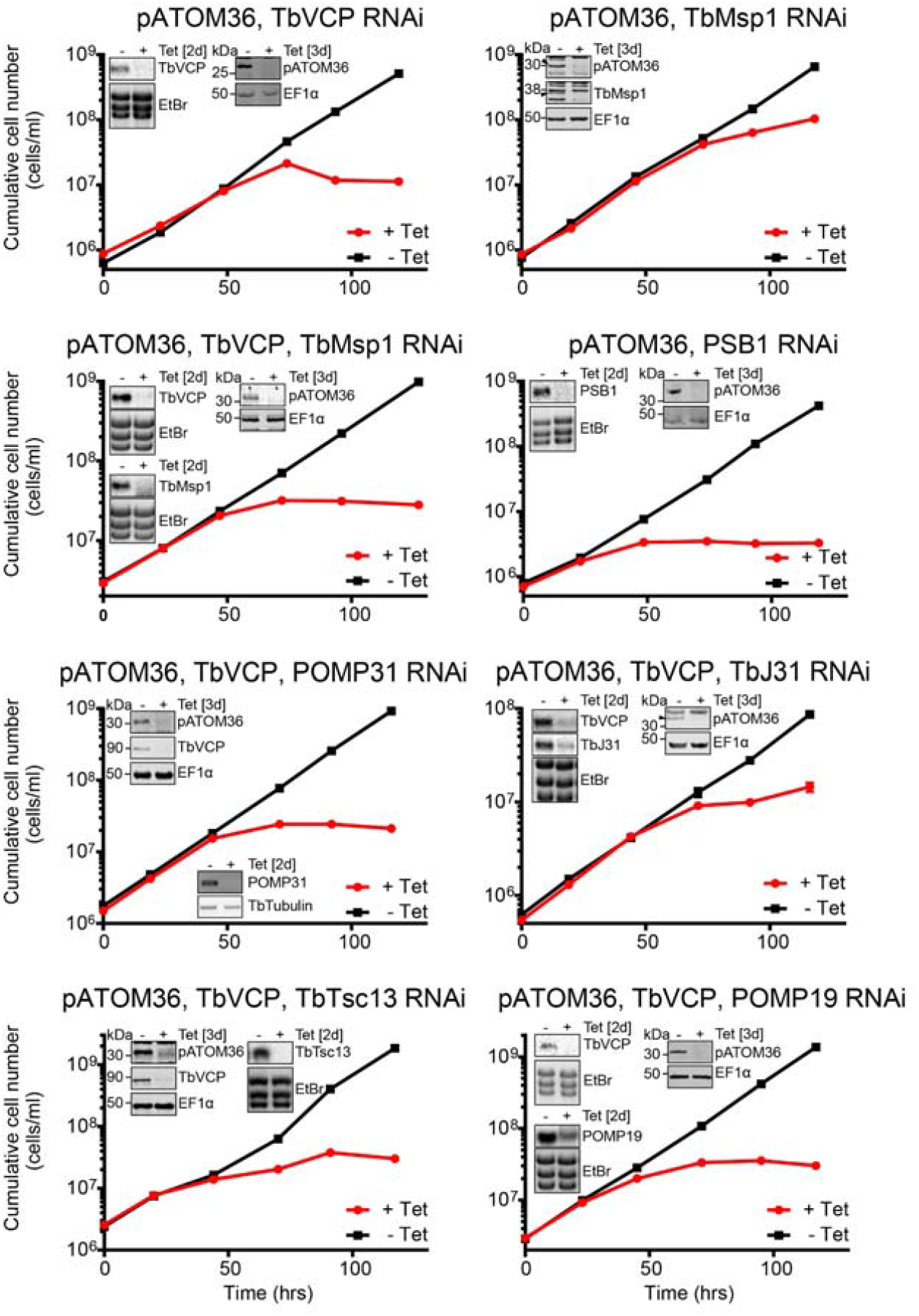
Verification and growth analysis of double and triple RNAi cell lines. Growth curve of induced (+Tet) and uninduced (-Tet) double and triple RNAi cell lines. Error bars correspond to the standard deviation (n = 3). The inset panels show the efficiency of RNAi for the indicated targets, either two days after induction when analysed by northern blots or RT PCR, or three days after induction when analysed by immunoblots. Ethidium bromide stained rRNAs, tubulin cDNA or EF1α serve as loading controls respectively.

## Notes

### Competing Interest Statement

The authors have declared no competing interest.

